# Human biliary atresia extrahepatic cholangiocyte organoids express increased ER and oxidative stress, altered drug metabolism and cell polarity changes

**DOI:** 10.1101/2025.05.04.649927

**Authors:** Adi Har-Zahav, Yara Hamoudi, Keren Danan, Ana Tobar, Michal Basphelchik, Michael Gurevich, Raanan Shamir, Irit Gat-Viks, Orith Waisbourd-Zinman

## Abstract

**Background & Aims:** Biliary atresia (BA), the leading cause of liver transplantation in children, presents in neonates with jaundice and progressive extrahepatic bile duct obstruction, yet its etiology and pathogenesis remain unknown. Here, we aimed to investigate the molecular mechanisms underlying BA and the susceptibility of cholangiocytes in the extrahepatic biliary tree using patient-derived extrahepatic cholangiocyte organoids (EHCOs).

**Methods:** EHCOs were derived from common bile ducts remnants of BA patients undergoing Kasai portoenterostomy and from non-BA controls at the time of liver transplantation. Transcriptomic profiling was performed via bulk RNA sequencing, and analyzed in two ways: differentially expressed pathways and perturbation analysis to predict aberrant functions. Key findings were validated through mechanistic assays, immunofluorescence staining, qPCR and transmission electron microscopy (TEM).

**Results:** Transcriptomic analysis predicted significant alteration in endoplasmic reticulum (ER) stress, dysregulations of drug metabolism, alongside pronounced alterations in cellular adhesion and polarity-related genes in BA-derived EHCOs. Cell-to-cell alterations were observed with various proteins including E-cadherin, RhoU, Sox17 and CFTR. BA EHCOs had an increased endoplasmic reticulum (ER) stress response, exemplified by elevated *PERK, BiP*, and *ATF4* along with abnormal ER on TEM. Furthermore *CHOP, ERO1A, WFS1*, and *SOD3* were decreased suggestive of abnormal ER stress response. BA EHCOs displayed increased toxicity to biliatresone-induced injury and inhibition of cytochrome P450 resulted in attenuation of the ER stress markers *PERK, BiP* and *ATF4*. Finally, liver hilum biopsies from BA patients undergoing Kasai portoenterostomy confirmed elevated PERK and PGR78(*BiP*) consistent with the EHCOs analysis.

**Conclusions:** BA EHCOs exhibit disrupted polarity, ER stress, and increased susceptibility to drug toxicity. These findings highlight key pathogenic mechanisms in BA and suggest that targeting these pathways may help mitigate cholangiocyte injury in BA.

**Impact and implications:** This study provides the first transcriptomic and functional analysis of human extrahepatic cholangiocyte organoids (EHCOs) derived from biliary atresia (BA) patients. By focusing on the extra-hepatic biliary tree, we identified key mechanisms of cholangiocyte injury, including persistent ER stress, impaired stress response pathways, altered drug metabolism and disrupted epithelial polarity. These findings highlight ER stress and metabolic vulnerability as potential therapeutic targets and establish EHCOs as a tractable model for investigating BA pathogenesis.

**Graphical abstract:** 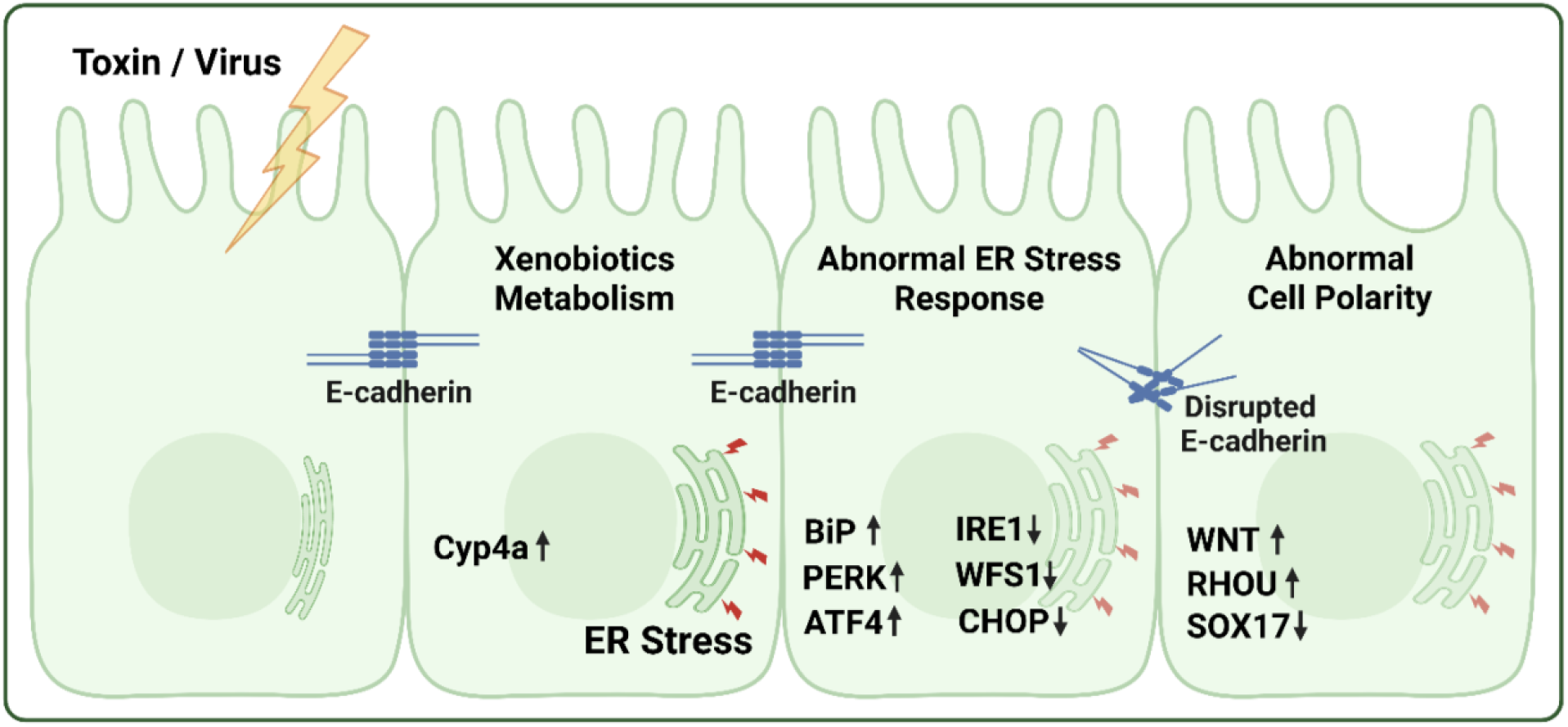

**Highlights:**

- First transcriptomic profiling of extrahepatic cholangiocyte organoids (EHCOs) from BA patients,
- revealing distinct molecular alterations compared to controls.
- BA EHCOs exhibit disrupted epithelial polarity, with downregulation of E-cadherin and Sox17 and upregulation of CFTR.
- ER stress is a hallmark of BA cholangiocytes, with elevated PERK, BiP, and ATF4, and
- dysregulation of downstream effectors including CHOP, ERO1A, and SOD3.
- BA EHCOs are more susceptible to biliatresone-induced injury, with enhanced ER stress and structural damage.
- Inhibition of cytochrome P450 activity (CYP4A) reduces ER stress markers.

## Introduction

Biliary atresia (BA) is a poorly understood cholangiopathy characterized by progressive extrahepatic bile duct obstruction in newborns. It is the leading cause of pediatric liver transplantation worldwide. Despite numerous years of research, the etiology and pathogenesis of BA remain elusive. The primary treatment is the Kasai portoenterostomy (KPE), a surgical attempt to restore bile flow from the liver to the intestine, however, outcomes remain unsatisfactory for many patients.^1^ A deeper understanding of the pathogenesis of BA, specifically understanding molecular mechanisms of extrahepatic bile duct injury in BA is critical for developing future therapeutic strategies.^2^

Clinically, the phenotype of BA at the time of presentation is more pronounced in the extrahepatic biliary tree compared to its intrahepatic counterpart. Even in cases where bile flow is successfully restored following KPE, patients often continue to experience progressive biliary fibrosis. BA is thus considered a pan-cholangiopathy, affecting both the intrahepatic and extrahepatic biliary systems. It is worth noting that the intra- and extrahepatic bile ducts originate from a different embryologic source. The extrahepatic ducts develop alongside the ventral pancreatic ductal system, later connecting with the intrahepatic ducts that originate from the hepatic bud.^3^ This distinction in embryological origin suggests that the molecular processes in the extrahepatic biliary tree during BA may differ from those in the intrahepatic bile ducts, necessitating further investigation, which is the focus of this study.

In recent years, transcriptomics has provided significant insights into the molecular landscape of BA. High-throughput RNA sequencing (RNA-seq) has identified differentially expressed genes, pathways, and potential molecular mechanisms underlying BA pathogenesis. Previous studies have highlighted immune-mediated damage,^4^ the interplay between fibrosis and the immune system,^5^ and abnormalities in ciliary function and epithelial cell development.^6^ While these insights have expanded our understanding of BA, they primarily reflect secondary consequences of bile duct obstruction rather than the initial pathogenic events. Given that BA manifests first in the extrahepatic biliary tree, where bile duct injury is most pronounced, it is essential to investigate this compartment specifically.

In this study, we used extrahepatic cholangiocyte organoids (EHCOs) derived from BA patients undergoing KPE and compared them to normal cholangiocyte organoids from patients undergoing liver transplantation for metabolic conditions. Through transcriptomic profiling, advanced analysis and functional validation, we aimed to investigate the mechanisms of injury in the extrahepatic biliary tree in BA. To our knowledge, this is the first study to compare the transcriptome of extrahepatic cholangiocyte organoids from BA patients with those from non-BA controls.

## Methods

### Organoids isolation and growth Primary biliary tissue

Primary extrahepatic biliary tissue was obtained from patients undergoing Kasai procedure, or liver transplantation, under an approved Institutional Review Board protocol (RMC 19-0072) (**Table S1**). The common bile duct was excised and placed immediately in UW solution for EHCOs generation.

### EHCOs generation and propagation

EHCOs were generated and propagated based on Tysoe OC et. al.^7^ Briefly, the common bile ducts were dissected and washed, and the luminal epithelium was scraped using a surgical blade while covered with William’s E media (Gibco). The cells were centrifuge at 500g for 4 minutes, washed and resuspended in a mixture of 66% Matrigel (Corning, 356237) and 33% William’s E medium (Gibco, Life Technologies) supplemented with 10 mM nicotinamide (Sigma-Aldrich),

17 mM sodium bicarbonate (Sigma-Aldrich), 0.2 mM 2-phospho-l-ascorbic acid trisodium salt (Sigma-Aldrich), 6.3 mM sodium pyruvate (Invitrogen), 14 mM glucose (Sigma-Aldrich), 20 mM HEPES (Invitrogen), ITS + premix (Corning, 354352), 0.1 μM dexamethasone (Sigma-Aldrich), 2 mM Glutamax (Invitrogen), 100 U/ml penicillin and 100 μg/ml streptomycin, 20 ng/ml EGF (PeproTech), 500 ng/ml R-spondin (PeproTech) and 100 ng/ml DKK-1 (PeproTech). 50µl of the cell suspension was plated in a 24-well plate so that a small dome of Matrigel formed in the center of each well. Plates were incubated at 37 °C for 2 minutes, then placed upside down and incubated for an additional 30 minutes until the Matrigel was solidified. Subsequently, 1 ml of William’s E medium with supplements was added to each well. The culture medium was replaced three times a week. Confluent wells were split at 1:4 ratio, and EHCOs were cryopreserved using NutriFreez D10 Medium (Biological Industries) at −80^0^c, then transferred into liquid nitrogen for long term preservation. All experiments were performed using EHCOs cultures that were previously cryopreserved and thawed in order to maintain consistent conditions. Organoid morphology was assessed using a brightfield microscopy and images were taken using ZEISS Primovert light microscope equipped with Axiocam 105 color camera, using x4 and x20 objectives.

### Bulk RNAseq

#### RNA extraction and RNA Bulk-Sequencing

RNA was extracted from 5 BA and 3 non-BA controls EHCOs at passage 2, using EZ-10 DNAaway RNA Miniprep Kit (BIO BASIC #BS88136) according to the manufacturer protocol. RNA integrity was analyze using TapeStation (Agilent Technologies), and samples (RIN>8) were sent to the Crown Genomics institute of the Nancy and Stephen Grand Israel National Center for Personalized Medicine (G-INCPM), Weizmann Institute of Science for library preparation and sequencing. The samples were sequenced using illumina NextSeq High Output - 75 cycles.

#### RNA-seq data of BA patients

The dataset consists of bulk RNA-seq profiles from EHCOs, originating both from patients with BA undergoing Kasai portoenterostomy (5 patients) and normal ducts from patients undergoing liver transplant for metabolic conditions (3 patients). For each of the eight samples, we used FASTQ files of the forward DNA strand (length 67bp). Quality control was performed using FastQC. As evidence for the quality of data, we observed the following metrics: on average, each sample contains 30.7M reads, with no reads flagged as poor quality, and the average GC content across samples is 51%.

Alignment of reads to the reference genome (hg38.refGene) was conducted using the STAR aligner, resulting in a reads-per-gene count matrix. Both the raw sequencing data and the count matrix were deposited in GEO (accession number GSE276230). The average mapping ratio was 83.54%, with a total of 28,271 genes detected. Gene filtration and normalization were performed using the DESeq2 R package, with default filtering settings and a minimum read count threshold of 10 reads per gene. After filtration, 16,896 genes were retained for subsequent analysis.

#### Analysis of differential expression in BA

Differential expression (DE) analysis of BA versus controls was performed through DeSeq2 in R (**Table S2**). For the log2FC value, positive/negative sign indicates up-regulation/down-regulation in BA compared to healthy samples. For each pathway (from KEGG and the GO repositories), we calculated the bias of the log2FC scores for genes within the pathway compared to the remaining genes. The bias is quantified using the Wilcoxon rank-sum test (FDR-adjusted *q* values). The ‘differentially expressed pathways’ are pathways with *q*-value < 0.005. Enriched pathways are further classified as either upregulated or downregulated pathways according to the direction of bias in BA – namely, ‘upregulated/downregulated pathways’ in BA are those that are enriched with genes that have high/low log2FC scores (**Table S3**).

#### Pre-processing of reference data

We used perturb-seq data, which targets genes with CRISPR interference (CRISPRi) with subsequent transcription phenotyping at single cell resolution.^8^ The experiment was performed in the retinal pigment epithelial (RPE1) cell line. We used as input the processed data after various filtration and pre-processing steps as reported in Replogle *et al*., 2022. The input data, which is used as input in our study, consists of a z-score for each gene in each single cell. The z-scores were obtained by normalization to control cells. The input data include 2393 perturbations, each of which was measured in 2 to 3461 single cells; each single cell is represented by a vector of z scores across 8748 transcribed genes. In addition, the input data includes non-targeting control cells.

For each genetic perturbation *r*, we applied the following two steps. First, we calculated the effect of the perturbation in gene *r* on each transcribed gene *g* by comparing the z scores of gene g in all single cells in which *r* is perturbed against all non-targeting (control) single cells (p-value of Wilcoxon rank-sum test, q-value after FDR correction for multiple genes). The ‘perturbation effect score’ of a perturbation *r* on a gene *g* is defined as the signed log10 of this *q*-value, with positive/negative perturbation effect scores for increase/decrease in the median values of gene *g* in *r*-perturbed cells compared to non-targeting cells. Each perturbation *r* is represented with a ‘perturbation profile’, consisting of a 8748-length list of the perturbation effect scores for perturbation *r* across all 8748 transcribed genes. A total of 2393 perturbation profiles were calculated, a profile for each perturbed gene. These profiles were used as input in the analysis of differential activation in BA.

#### Analysis of differential activation in BA

The analysis takes as input: (1) A BA gene sets: either a BA-upregulation gene set, including the 100 genes with highest log2FC scores (denoted BA-Up), or the BA-downregulation gene set, including the 100 genes with lowest log2FC scores (denoted BA-Down) (**Table S2**). (2) Perturbation effect scores of a given factor. This reference data consists of the perturbation profiles of a factor across the 8748 transcribed genes. Of note, only 18 of the 100 BA-Up genes and only 10 of the 100 BA-Down genes were expressed in perturbed single cells and were therefore used in the subsequent calculation.

Given an input BA gene set (BA-Up or BA-Down sets) and a perturbation profile of a given factor, the ‘effect of perturbation on a BA gene set’ is defined as the bias of the perturbation effect scores of the perturbed factor on genes within the BA gene set compared to the remaining genes (a Wilcoxon rank-sum test *p* value) (**Table S4**). Thus, a perturbation upregulates/downregulates the BA gene set when the genes in the BA gene set tend to high/low perturbation effect scores. We distinguish four types of factors with a significant effect (**Table S4**): Factors whose perturbation downregulates the BA-Down’s genes (25 factors, effect on BA-Down *p* <0.11); (2) downregulates the BA-Up’s genes (48 factors, on BA-Up p<0.01); (3) upregulates the BA-Down’s genes (25 factors, effect on BA-Down p< 0.07); and (4) upregulates the BA-Up’s genes (2 factors, effect on BA-Up p<0.08). For the factors within each of these categories, we performed hyper-geometric test using each of the pathways of the REACTOME collections (**Table S5**). Pathways that are enriched (*q* <0.05) are referred to as ‘differentially activated pathways’.

#### Immunofluorescent (IF) stains and microscopy

EHCOs were plated on 15 well and 18 well µ-slide with glass bottom (ibidi #81817, #81507) in 10µl mixture of 2:1 Matrigel and E+ supplemented media, covered after solidifying with additional 50 µl supplemented media. Media was replaced 3 times a week for 1-2 weeks, and the EHCOs were fixed using 4% PFA for 20 minutes in 4^0^c, followed by 3 washes with 1xPBS. EHCOs were incubated with sera containing permeabilization solution (10% FCS, 1% BSA, 2% gout serum, 0.2% TTX) for 1 hour at room temperature, then antibodies were added and incubated over-night at 4^0^c, followed by three washes with permeabilization solution (45 minutes each, at room temperature). Secondary antibodies were added together with Phalloidin and incubated overnight at 4^0^c. The next day the slides were washed 3 times with PBSx1, stained with DAPI for 10 minutes at room temperature, washed again and covered with PBSx1. Images were generated using Leica SP8 confocal microscope, at x20 and x40 magnification, images were analyzed using ImageJ software. In order to avoid bias, the IF staining pictures were taken from normal appearing EHBCs.

#### Real time quantitative PCR

EHCOs were harvested using Cell Recovery Solution (Corning) and RNA was extracted using RNeasy Mini Kit (QIAGEN). cDNA was prepared using a qScript TM cDNA Synthesis kit (Quanta bio). Quantitative PCR was preformed using StepOne Plus and SYBR green reagent (applied Biosystems). A list of primers can be found in **Table S6**. Expression analysis was done using the double delta ct method as previously described.^9^

#### ER immunofluorescence stain

EHCOs were dissociated into single cells and plated as monolayer: EHCOs were harvested using Cell Recovery Solution (Corning) and placed on ice for 30 minutes, then washed and incubated in 1ml TrypLE Express Enzyme (Gibco) for 5 minutes in 37^0^c. Followed pipetting another 1ml supplemented William’s E media was added and the cells were stained using pre-washed 30µm cell strainer (PluriSelect). 100,000 cells per well were seeded on a Lab-Tek II 8 well plate (Nunc) covered with 1mg/ml rat tail collagen (adjusted from.^10^ Cells were fixed with 3.6% PFA and stained using the ER-ID green assay kit (ENZO #51025) according to the manufacturer’s protocol. Images were generated using Nikon AX confocal microscope at x60 magnification and processed using ImageJ software.

#### TEM microscopy

EHCOs were harvested and placed in Karnovski (Glutarhaldehide 70%, Formalin 37%, Cacodylate buffer0.2M,H_2_O), then transferred to Rabin medical center TEM pathology core facility to prepare for microscopic imaging. Images were taken using JEM-1400Flash microscope.

#### HET0116 treatment

EHCOs were treated with 8ug/ml Cytochrome P450 inhibitor, HET-0016 (CAS 339068-25-6, Santa Cruz) for 6 hours. EHCOs were harvested, and RNA was extracted. ER stress related genes expression was measured by RT-qPCR and compared to untreated EHCOs.

#### Biliatresone treatment

EHCOs cultures were treated with either 5 µg/mL or 10 µg/mL biliatresone (HY-119412, MCE) for 72 hours. An equivalent volume of DMSO was used as a vehicle control. The effect of biliatresone was assessed by evaluating EHCOs integrity or performing IF stain. To assess organoid integrity, brightfield images were acquired 72 hours post-treatment using a ZEISS Primovert light microscope equipped with an Axiocam 105 color camera and a 4× objective. EHCOs were classified as either intact or deformed and quantified using ImageJ software. For each condition, at least 35 EHCOs were analyzed per culture across three independent experiments. For IF staining, EHCOs were fixed with 4% PFA and processed as described in the “IF Stains and Microscopy” section. This analysis was performed in five independent experiments.

#### Staining of paraffin-embedded human slides

Paraffin-embedded slides from the liver hilum obtained from BA patients and non-BA controls were stained for PERK and BiP antibodies (**Table S7**). Briefly, slides were deparaffinized by warming to 60 °C, treated with xylene, then rehydrated with decreasing concentrations of ethanol (100%, 95%, 80%, 70%). Slides were incubated in microwave oven for 15 min in citric acid buffer (pH 6), cooled, and washed in running water, washed with PBS, blocked with PBT, and cultured with primary antibodies diluted in PBT over-night at 4 °C, washed with PBT, incubated with secondary antibodies for 30 min in RT, washed two times, stained with DAPI (5 min, RT), washed, and covered with mounting medium and a cover slip. Images were taken using Nikon AX Confocal microscope at x40 magnification and analyzed using ImageJ.

#### Statistical analysis

For the experiments other than the RNA-seq analysis, unless stated otherwise, mean, standard error, and p values were calculated using GraphPad Prism 9. Statistical significance was calculated by unpaired Student’s t test, p value <0.05 was considered statistically significant.

## Results

### BA extrahepatic cholangiocyte organoids differ from normal controls in morphology and gene expression

We generated extrahepatic human cholangiocyte organoids (EHCOs) from common bile duct remnants of BA patients and controls and grew them as organoids (**Figure 1A,B**). Patients’ derived human organoids clinical data are presented in **Table S1**. Successful propagation rate of BA derived EHCOs was 65%, while 91% of the control EHCOs were propagated into viable organoid cultures. Moreover 6 out of 15 (40%) BA derived EHCOs cultures demonstrated deformed organoid shape, compared to 1 deformed EHCOs out of 11 controls (9%) (**Figure 1B,C**). EHCOs sizes were not uniform between patients and had a wide range in each culture, of approximate 50-500 microns, regardless of the baseline patients’ characteristics. In order to study mechanisms of cholangiocyte injury of the extrahepatic biliary tree of patients with BA we performed RNA-seq from BA and control organoids. We profiled RNA from 5 BA patients and 3 controls and used passage 3 culture for all samples.

**Figure 1:**
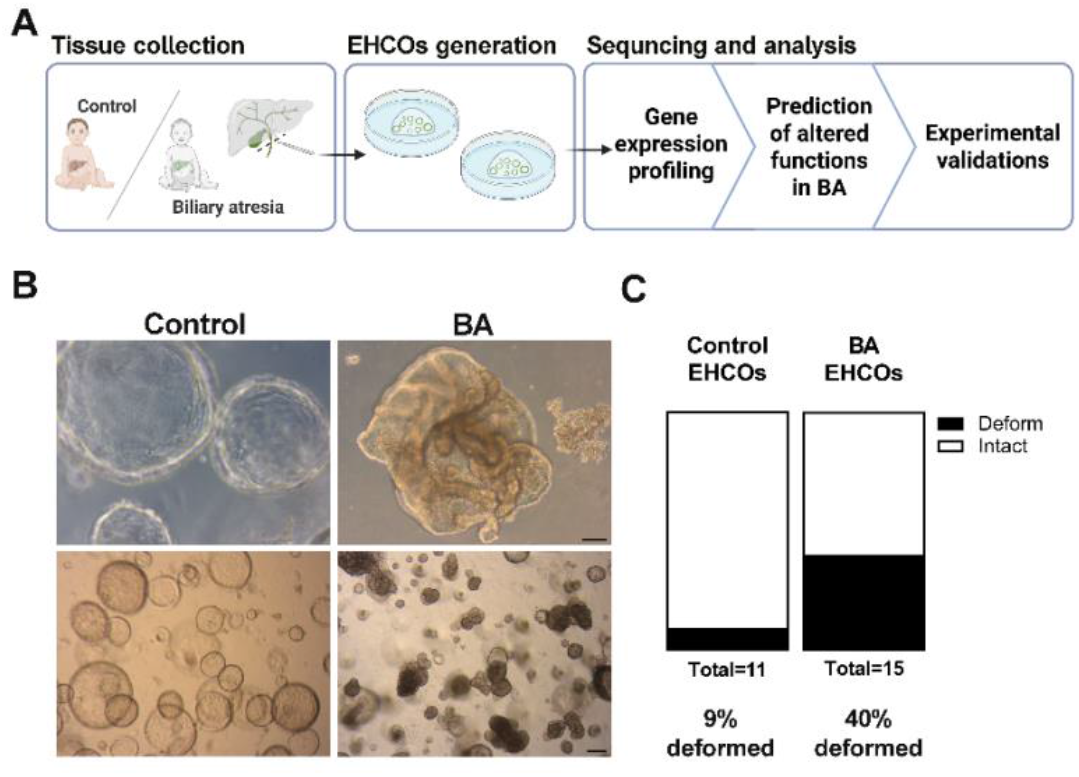
Study design and cholangiocyte organoids propagation. (a) Study design: extrahepatic bile ducts were obtained from BA patient and non-BA controls and propagated as human cholangiocyte organoids (EHCOs). EHCOs underwent bulk RNA-seq and transcriptomic analysis. (b) Representative images of intact control EHCOs, and deformed BA EHCOs. Upper panel scale bar=200 um, lower panel scale bar=100 um. (c) Quantification of EHCOs morphology as open lumens or deformed shapes in BA (n=15) and non-BA controls (n=11).

Estimates of the BA-associated change in gene expression revealed a robust response, including both up- and down-regulation of genes (**Figure 2A, Table S2, Methods**). **Figure 2B** highlights the genes with the top fold changes, including the top 100 upregulated and top 100 downregulated genes. Gene set enrichment analysis (elaborated under the methods section) of the up/down-regulated genes showed enriched upregulation of genes in drug-metabolism pathways, such as ‘drug metabolism by cytochrome P450’ (enrichment *q* <10^−5^), ‘metabolism of xenobiotics by cytochrome P450’ (*q* <10^−5^), as well as upregulation of pathways related to epithelial integrity, such as ‘biological adhesion’ (*q* <10^−4^) and ‘cell junction organization’ (*q* <10^−3^). We also observed enriched downregulation of genes in translation (e.g., ‘ribosome’ *q* <10^−21^), oxidative phosphorylation (*q* <10^−6^) and cell cycle (q <10^−3^) processes (**Table S3**). Next, we analyzed the DE genes in conjugation with a precompiled dataset of gene expression changes induced by various perturbations (elaborated under the methods section). This analysis enabled the prediction of factors that have a significant effect on up/down-regulated genes, and subsequently the prediction of pathways that are enriched with these factors (**Table S4**,**S5, Methods**).

**Figure 2:**
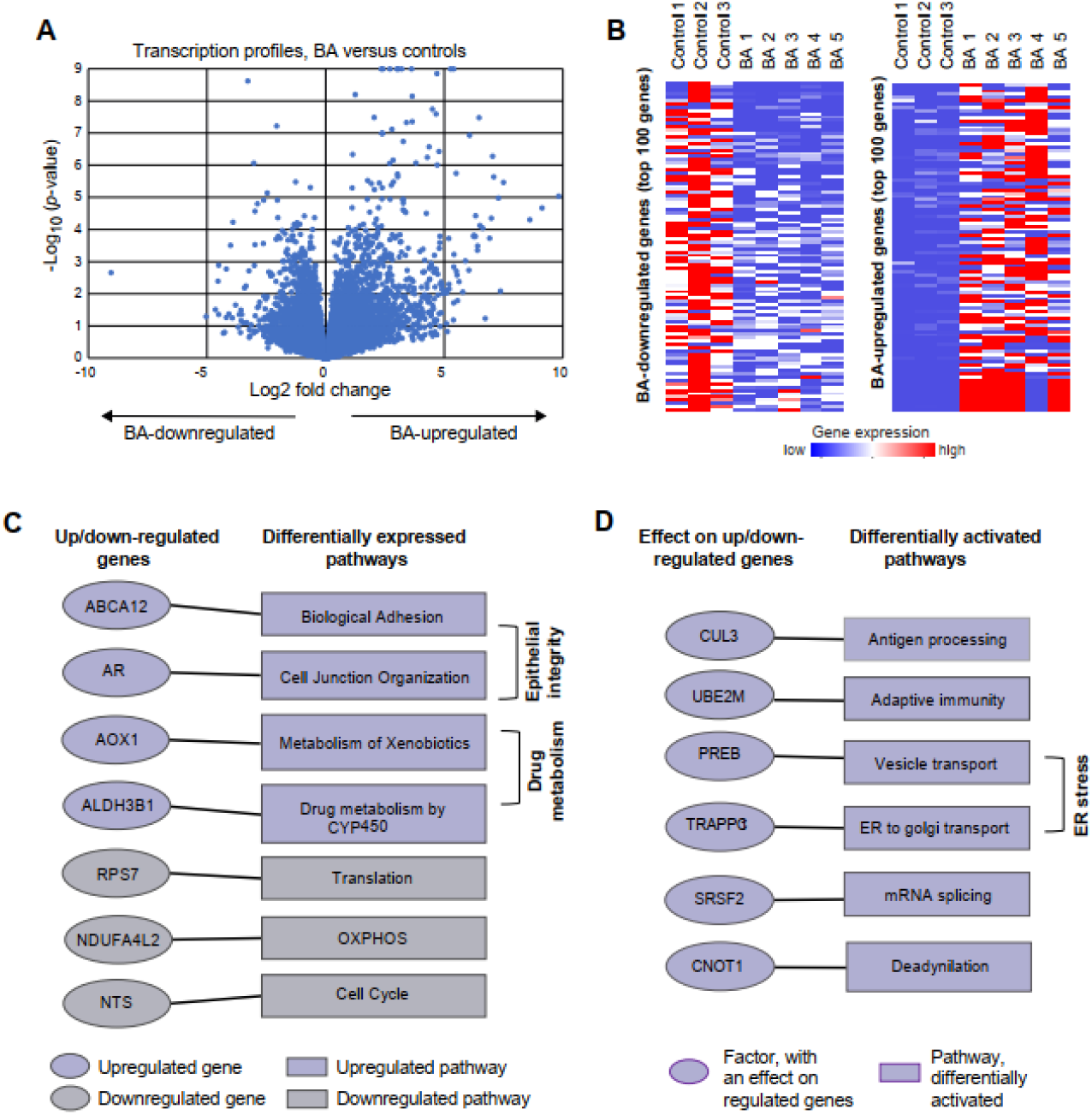
Molecular profiling reveals transcriptomic changes in BA HCOs. (a) Differential expression between BA compared to controls. Volcano plot showing the comparison of transcription profiles between EHCOs derived from 5 BA patients and 3 non-BA controls. (b) Top up- and down-regulated genes. A heatmap presenting the expression level (color coded) of each gene (columns) in each individual (rows). Five BA patients and three controls are indicated. Included are top 100 genes that are up-regulated (left) or up-regulated (right) in BA. (c) Differentially expressed pathways. Left: examples of up/down-regulated genes (color coded; see full list in Table S2). Right: examples of pathways enriched with upregulated or downregulated genes (color coded; see full list in Table S3). Genes and their associated pathways are connected by edges. (d) Differentially activated pathways. Left: examples of factors that have an effect on up/down-regulated genes in BA (see full list in Table S4). Right: examples of pathways enriched with the identified factors. The identified pathways are referred to as “differentially activated pathways” (see full list in Table S5). Factors and their associated pathways are connected by edges.

Pathways that are enriched with the identified factors, referred to as ‘differentially activated pathways’ include vesicle transport, such as ‘ER to Golgi transport’ (*q* <10^−4^, e.g., TRAPPC1), and metabolism of RNA, such as ‘mRNA splicing (*q* <10^−7^, e.g., SRSF2((**Figure 2D, Table S5**). Overall, based on these predictions, we set to focus on three aspects: epithelial integrity, ER stress and drug metabolism (**Figure 2C,D**). We performed further experiments to validate the relevance of these hypothesis. The validation and functional studies were done with additional EHCOs samples from both BA and control groups (**Table S1**).

### BA EHCOs exhibit aberrant cell polarity and epithelial function

Cell-to-cell adhesion and cell polarity are important for the proper epithelial function and regulate tissue homeostasis in critical cell processes that include tissue barrier function, cell proliferation, and migration.^11^ This is particularly relevant for the biliary tree epithelium in which improper barrier function may result in bile leak and exposure of cholangiocytes and the sub epithelial layer to bile. E-cadherin is a calcium-dependent adhesion molecule that helps maintain the integrity and polarity of cholangiocytes. It is a key component of adherents junctions and essential to maintain the polarity of the biliary epithelium.^12,13^ EHCOs form BA patients had decreased E-cadherin IF expression compared to normal EHCOs (n=5, p=0.0430) (**Figure 3A**). RhoU, also known as Wrch1, is an atypical member of the Rho GTPase family that plays a crucial role in regulating cell polarity, cytoskeletal dynamics, and cell adhesion).^14,15^ RhoU IF stain was increased in BA EHOCs compared to controls (n=5, p<0.0001) (**Figure 3B**). SOX17 is a transcription factor known to be important in cell polarity,^16–18^ EHCOs from BA patients had decreased expression compared to controls (n=5, p=0.0167) (**Figure 3C**). The cystic fibrosis transmembrane conductance regulator (CFTR) protein is a cAMP regulated chloride and bicarbonate ion channel expressed at the apical plasma membrane of the extrahepatic biliary epithelium which has a significant role in in cell junction formation and actin cytoskeleton organization with its connection to the ECM.^19,20^ Interestingly BA EHCOs overexpress CFTR compared to normal control (n=5, p=0.0005) (**Figure 3D**). This potentially can be a compensatory mechanism to restore epithelial organization as well as improve poor bile flow via decreased bile viscosity. Overall, these data demonstrate changes in epithelial integrity and cell junction organization in BA EHCOs compared to controls.

**Figure 3:**
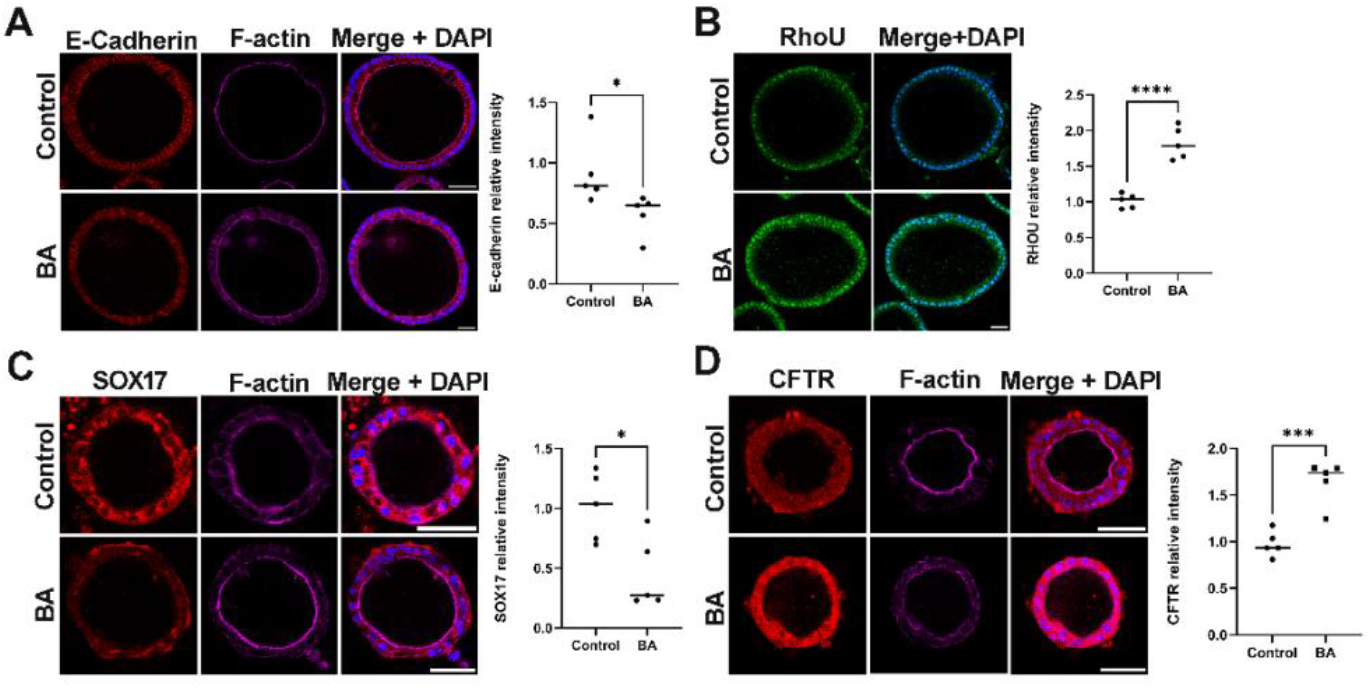
BA-derived EHCOs demonstrate disruption of epithelial polarity. (a) IF stains for E-cadherin (red), F-actin (magenta), and DAPI (blue) in EHCOs derived from BA and non-BA controls, n=5 in each group, p=0.0430, scale bar= 50 μm. (b) IF stains for RhoU (green) and DAPI (blue) in EHCOs derived from BA and non-BA controls, n=5 in each group, p< 0.0001, scale bar= 50 μm. (c) IF stains for Sox17 (red), F-actin (magenta), and DAPI (blue) in EHCOs derived from BA and non-BA controls, n=5 in each group, p=0.0167, scale bar= 50 μm. (d) IF stains for CFTR (red), F-actin (magenta), and DAPI (blue) in EHCOs derived from BA and non-BA controls, n=5 in each group, p=0.0005, scale bar= 50 μm.

### BA EHCOs demonstrate increased ER stress with abnormal stress response

The differential regulation analysis revealed changes in functions of vesicle transport and mRNA splicing in ER in BA EHCOs. The ER is essential for synthesizing, modifying, and folding proteins. Moreover, ER stress can activate the non-canonical WNT signaling pathway leading to disruption of cytoskeleton organization, which plays a role in cell polarity.^21^ Based on these findings, we investigated ER stress and the associated stress response pathways in BA. We first stained human BA cholangiocytes for the ER IF marker ER-ID,^22^ which showed higher stain along with condensed ER in BA cholangiocytes compared to controls suggestive of ER stress (n=4 in each group) (**Figure 4A**). We also looked at the ER morphology of EHCOs via TEM which revealed dilated and expanded ER folds also suggestive of ER stress^23,24^ (n=3 in each group) (**Figure 4B**). We then performed qRT-PCR for several ER stress markers. BiP (Binding immunoglobulin protein, also known as GRP78,encoded by HSPA5) is a chaperone protein that can switch from its normal chaperone function to become an ER stress sensor,^25^ and found it was upregulated in BA HCOs compared to controls (n=8 in BA, 5 in non-BA controls, p=0.0328) (**Figure 4C and Figure 7**). Under normal conditions, BiP binds to the luminal domains of the UPR (unfolded protein response) sensors (IRE1, PERK and ATF6) keeping them inactive. During ER stress BiP dissociates from these sensors to bind misfolded proteins, triggering UPR activation. We thus performed qRT-PCR to those three and found *PERK* (Protein kinase R-like ER kinase, encoded by *EIF2AK3*), and its downstream transcription factor ATF4^26^ were both upregulated in BA HCOs (n=5-7 in BA,5-6 in non-BA controls, p=0.0327, p=0.0073 respectively)(**Figure 4C and Figure 7**).IRE1alpha (Inositol-Requiring Enzyme 1 alpha, encoded by *ERN1*) a key transmembrane protein involved in the ER stress response that regulates various cellular processes related to cell survival and death.^27^ Interestingly, while it may have been speculated that *ERN1* (IRE1) would increase in a similar manner to *PERK. ERN1* gene expression was decreased in BA EHCOs compared to control (n=6 in BA, 5 in non-BA controls, p=0.0034) (**Figure 4C and Figure 7**). ATF6 (Activating Transcription Factor 6) plays a pivotal role in linking the ER stress response to oxidative stress, contributing to cellular adaptation and survival under stress conditions.^28–31^ Thus, it was expected to increase with ER stress, however the change in *ATF6* expression was not significantly different between BA and control samples (n=6 in BA, 5 in non-BA controls, p=0.0611) (**Figure 4C and Figure 7)**. The decrease of *ERN1* (IRE1) and the lack of increase in *ATF6* may occur due to abnormal cholangiocyte response to ER stress or chronic ER stress. IF stain for BiP and PERK also showed increased stain in BA EHCOs compared to controls (n=5 BA, 5 control, p=0.0478, p=0.0004 respectively) (**Figure 4D, E**). We next examined the downstream effectors of PERK. Wolframin ER Transmembrane Glycoprotein (*WFS1*), *DDIT3* (encodes for C/EBP Homologous Protein (CHOP)) Superoxide Dismutase 3 (*SOD3*) and Endoplasmic Reticulum Oxidoreductase 1 Alpha (*ERO1alpha*), were all decreased in BA EHCOs compared to controls (n=5-6 BA, 3-5 control, p=0.0087, p=0.0004, p=0.0062, p=0.0284 respectively) (**Figure 4C**). While these are typically upregulated during prolonged ER stress and function to alleviate cell stress^32,33^ their decrease may suggest abnormal ER stress response. In addition, BA derived EHCOs demonstrated lower levels of reduced glutathione (*GSH*) stain compared to non-BA controls (n=5 BA, 5 control, p=0.0355) (**Figure 4F**).

**Figure 4:**
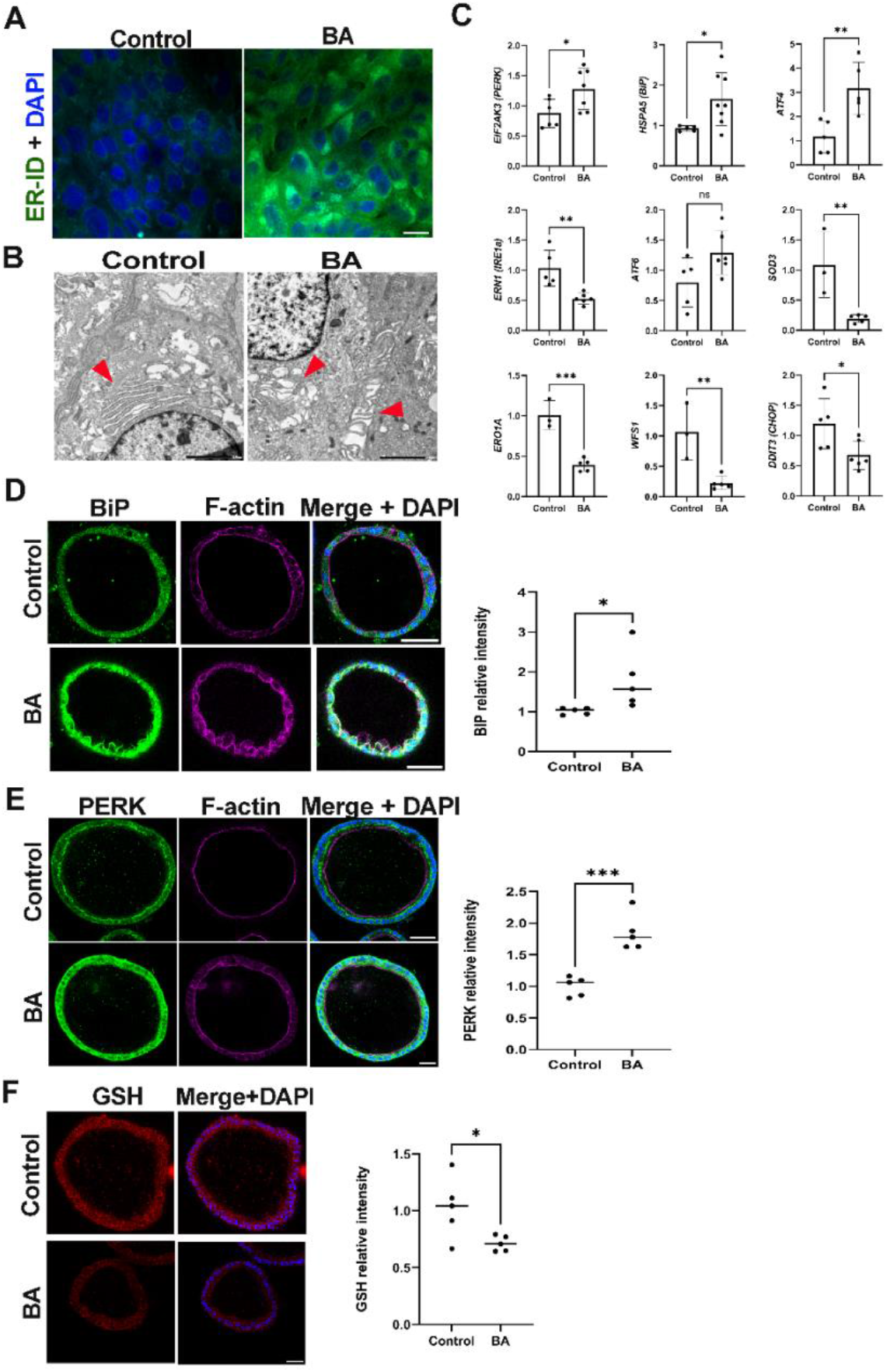
BA-derived ECHOs exhibit increased endoplasmic reticulum (ER) stress. (a) ER-ID staining of cholangiocytes from BA patients and non-BA controls, n=4 in each group, scale bar= 20μm. (b) TEM images of EHCOs derived from BA patients and non-BA controls. ER is indicated by red arrowheads, n=3 in each group; scale bar= 2 um. (c) qRT-PCR analysis of HCOs from BA patients and controls showing expression for *EIF2AK3, HSPA5, ATF4, SOD3, ERO1A, WFS1, DDIT3, ERN1* and *ATF6*. n=5-8 in BA, 3-6 in non-BA controls, p=0.0327, p=0.0328, p=0.0073, p=0.0087, p=0.0004, p=0.0062, p=0.0284, p=0.0034 and p=0.0611 respectively. (d) IF stains for BIP (green), F-actin (magenta), and DAPI (blue) in HCOs from BA and non-BA controls, n=5 in each group, p=0.0478, scale bar= 50 μm. (e) IF stains for PERK (green), F-actin (magenta), and DAPI (blue) in EHCOs from BA and non-BA controls, n=5 in each group, p=0.0004, scale bar= 50 μm. (f) IF stains for GSH (red), and DAPI (blue) in HCOs from BA and non-BA controls, n=5 in each group, p= 0.0355, scale bar= 50 μm.

### BA EHCOs have increased susceptibility to drug/toxic injury which augments stress response

Drug metabolism and cytochrome p450 were noted in the RNAseq analysis to differ between BA and control EHCOs. In order to determine response to drug injury in the EHCOs we used the isoflavonoid toxin biliatresone.^17,18,34^ We previously showed the biliatresone causes lumen obstruction of cholangiocyte spheroids obtained from the extrahepatic cholangiocytes of balb/c mice.^17,35^ We were thus intrigued to determine biliatresone effect on EHCOs and to determine if BA EHCOs organoids have increased susceptibility to biliatresone. We counted open EHCOs vs. obstructed ones in culture and compared this ration pre- and post-biliatresone treatment. Biliatresone resulted in significantly more obstructed EHCOs in BA compared to control EHCOs (n=3, p=0.0149, at least 35 EHCOs were counted for each sample) (**Figure 5A**). We then measured biliatresone-induced ER stress by staining the EHCOs for PERK and E-cadherin. Biliatresone treated BA EHCOs had a higher increase in PERK IF stain (n=5 BA,5 control, p=0.0103 (**Figure 5B**). E-cadherin IF stain showed an irregular and fragmented pattern (**Figure 5B**). We then inhibited cytochrome P4504A activity via the compound HET0016 and measured the previously shown ER stress markers via qPCR. *HSPA5, ATF4* and *EIF2AK3* were all decreased with HET0016 treatment (n=5, p=0.0034, p=0.0417, p=0.0007 respectively) (**Figure 5C**). Lastly, we measured *CYP4A11* via qPCR and found that BA EHCOs have increased expression compared to controls (n=5 BA,3 control, p=0.0399) (**Figure 5D**).

**Figure 5:**
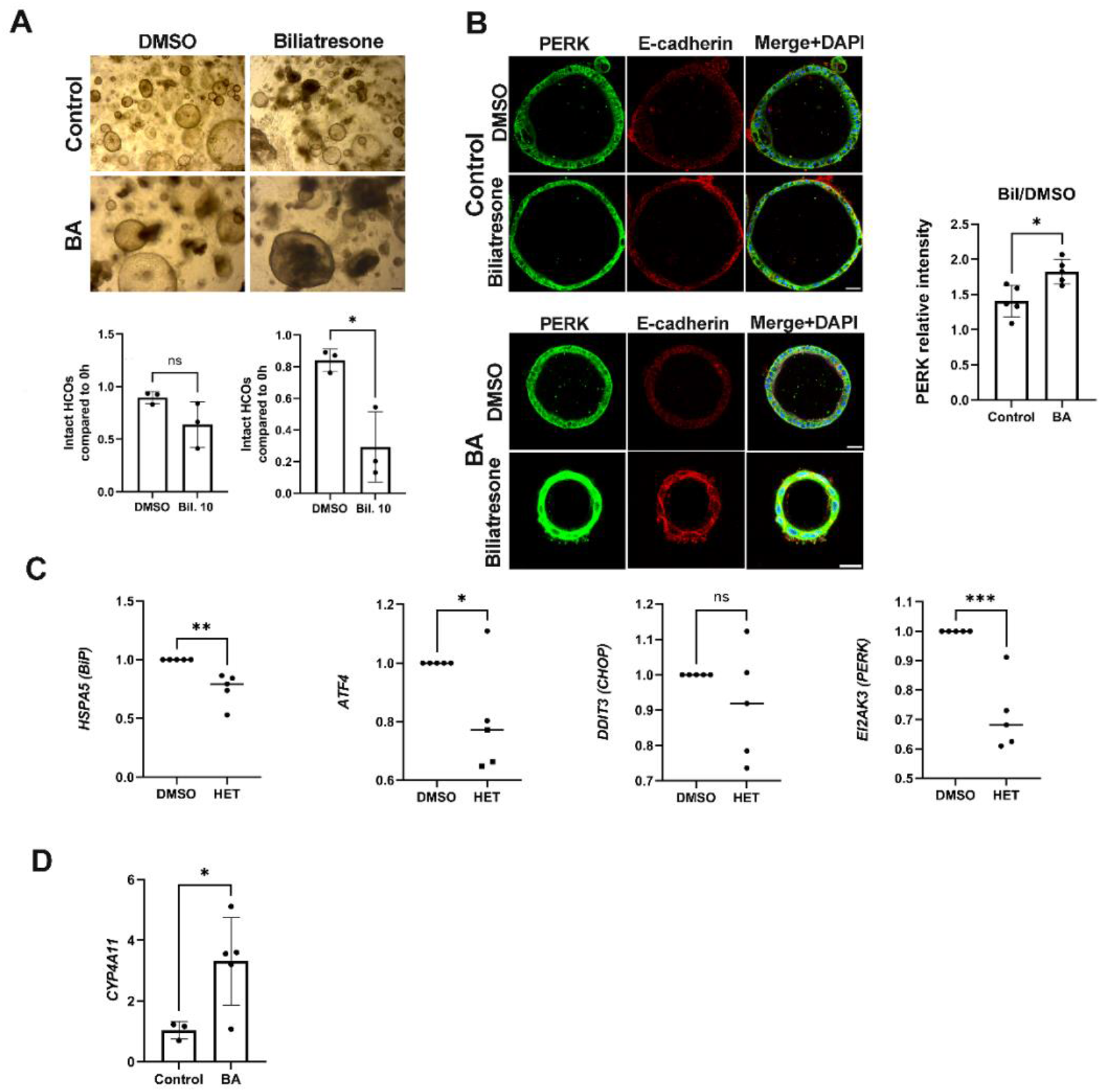
BA HCOs show heightened drug-induced endoplasmic reticulum stress. (a) Brightfield images of HCOs from BA and non-BA controls treated with biliatresone, along with quantification EHCOs by morphology (intact lumens vs. obstructed), n=3, scale bar= 200 μm, at least 35 EHCOs were counted per condition. (b) IF stains for PERK (green), E-cadherin (red), and DAPI (blue) in EHCOs from BA and non-BA controls treated with biliatresone or DMSO, n=5 for each group, p=0.0103; scale bar in control HCOs= 50 μm, scale bar for BA HCOs upper panel=50 μm, lower panel=30 μm. (c) qRT-PCR results of ER stress markers in BA EHCOs treated with HET0016. n=5 in each group, p=0.0034 (HSPA5), p=0.0417 (ATF4), p=0.0007 (EIF2AK3). (d) qRT-PCR expression of CYP4A11 in HCOs from BA and non-BA controls, n=5 BA; 3 non-BA, p= 0.0399. e bar= 200 μm, at least 35 EHCOs were counted per condition. (b) IF stains for PERK (green), E-cadherin (red), and DAPI (blue) in EHCOs from BA and non-BA controls treated with biliatresone or DMSO, n=5 for each group, p=0.0103; scale bar in control HCOs= 50 μm, scale bar for BA HCOs upper panel=50 μm, lower panel=30 μm. (c) qRT-PCR results of ER stress markers in BA EHCOs treated with HET0016. n=5 in each group, p=0.0034 (HSPA5), p=0.0417 (ATF4), p=0.0007 (EIF2AK3). (d) qRT-PCR expression of CYP4A11 in HCOs from BA and non-BA controls, n=5 BA; 3 non-BA, p= 0.0399.

### Human BA liver hilum biopsies exhibit increased ER stress proteins

Lastly, though EHCOs were obtained from human extrahepatic bile ducts and it was previously shown that EHCOs maintain their characteristics in culture.^7^ We wanted to correlate our findings with human biopsies. We stained liver hilum biopsies from BA patients (removed at the time of KPE) and controls (taken from the liver hilum at the time of liver transplant) and stained for the ER stress markers BiP and PERK. Both were increased in BA samples (n=3 BA, 4 control, p=0.0434, p=0.0433 respectively) (**Figure 6**).

**Figure 6:**
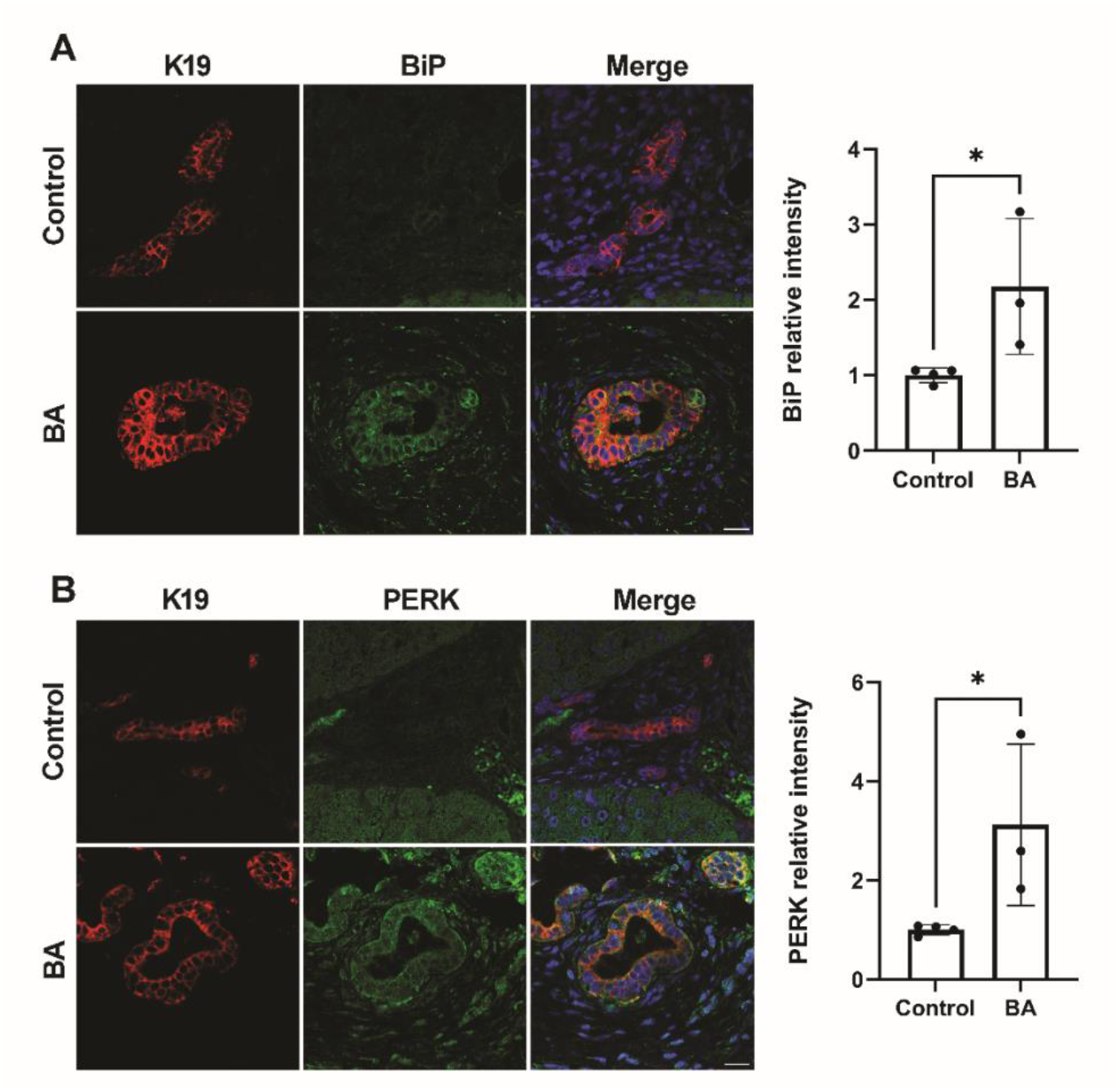
PERK and Bip are overexpressed in liver hilum biopsies from BA patients. Liver hilum biopsies from BA patients (collected at the time of Kasai portoenterostomy) and non-BA controls, (collected during liver transplantation) were stained for ER stress markers BiP and PERK. (a) IF stains of BiP (green), K19 (red), and DAPI (blue), n=3 for BA, n=4 for non-BA control, p=0.434, scale bar=50 μm. (b) IF stains of PERK (green), K19 (red), and DAPI (blue). n=3 for BA, n=4 for non-BA controls, p=0.04330, scale bar=50 μm.

**Figure 7:**
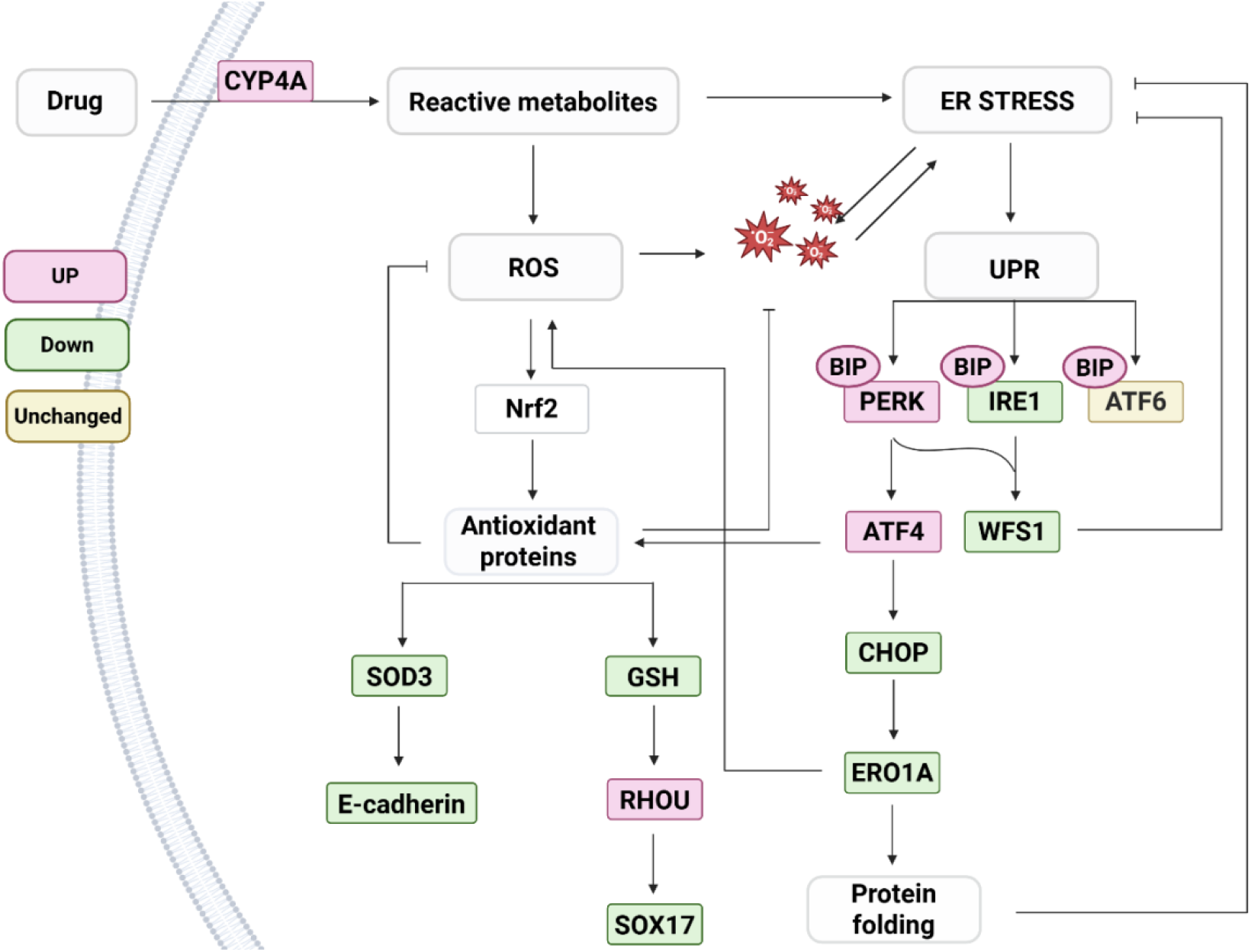
Pathways involved in BA EHCOs mechanism of injury. Summary of genes and pathways that were identified on bulk RNA-seq and further verified between BA and control HCOs. Created with Bio Render.

## Discussion

In this study, we conducted a comprehensive transcriptomic and functional analysis of extrahepatic cholangiocytes derived from patients with BA and compared them to controls. Our findings revealed key molecular and cellular alterations in the extrahepatic biliary epithelium, highlighting disrupted epithelial integrity, elevated endoplasmic reticulum (ER) stress, and altered drug metabolism as contributors to cholangiocyte susceptibility and injury. Unlike previous studies that focused on intrahepatic cholangiocyte organoids or whole liver transcriptomics, our work specifically examined the extrahepatic biliary tree. This distinction is crucial, as BA manifests primarily as an obstructive disease of the extrahepatic ducts, which have a distinct embryological origin from intrahepatic bile ducts. Additionally, while cholangiocytes can be derived from induced pluripotent stem (iPS) cells, these cells resemble fetal intrahepatic cholangiocytes. In contrast, our organoid model more accurately recapitulates the characteristics of extrahepatic cholangiocytes, as previously demonstrated.^7^ This approach provides a unique platform to elucidate the pathophysiological mechanisms underlying BA.

The transcriptomic analysis and subsequent validation experiments reveal significant alterations in pathways related to cell junction organization and polarity in BA-derived EHCOs. Specifically, we observed reduced expression of E-cadherin and Sox17, both essential for maintaining cholangiocyte polarity and epithelial integrity, alongside an increase in RhoU, a regulator of cytoskeletal dynamics and adhesion.^11,12,20^ These findings align with our previous work on extrahepatic cholangiocyte injury in a toxic model of biliary atresia, where exposure to biliatresone led to tight junction disruption and increased epithelial permeability, implicating genes in the non-canonical WNT, Notch, and Hippo signaling pathways.^17,18,34^ Supporting this, Amarachintha et al. demonstrated that biliary organoids derived from liver biopsies of BA patients exhibit polarity defects and reduced tight junction integrity in both intrahepatic BA organoids and liver tissues.^36^ Additionally, genome-wide sequencing analysis by Glessner et al. identified single nucleotide polymorphisms in *AFAP1* and *TUSC3*, key regulators of ciliogenesis and planar polarity, further underscoring the role of disrupted epithelial integrity in BA pathogenesis.^6^

Interestingly, CFTR was overexpressed in BA-derived EHCOs. CFTR is a cAMP-regulated chloride (Cl−) and bicarbonate (HCO3−) ion channel expressed at the apical membrane of cholangiocytes. Beyond its ion transport function, CFTR plays a crucial role in establishing and maintaining epithelial apical-basolateral polarity.^19^ Roos et al. demonstrated that CFTR activity is suppressed under hypoxic conditions in intrahepatic cholangiocyte organoids.^37^ In our study, CFTR overexpression may represent an adaptive or compensatory response to impaired epithelial architecture or an attempt by extrahepatic cholangiocytes to enhance bicarbonate secretion and reduce bile viscosity. Notably, Amarachintha et al. reported decreased CFTR expression in BA-derived organoids, though these originated from intrahepatic cholangiocytes.^36^ Collectively, these findings suggest that disruptions in cholangiocyte polarity within the extrahepatic biliary tree may increase epithelial permeability and compromise barrier function, potentially exacerbating bile duct injury and accelerating disease progression.

A key finding in this study was the evidence of ER stress in BA-derived EHCOs, supported by the upregulation of ER stress markers and morphological changes observed via electron microscopy. BiP, a molecular chaperone that alleviates cellular stress by binding misfolded proteins in the ER,^38^ and PERK, a key sensor of the UPR that reduces protein synthesis under stress conditions,^39^ were both significantly increased in BA-derived EHCOs. Additionally, ATF4, a transcription factor downstream of PERK that regulates restoration of cellular homeostasis after oxidative stress,^40,41^ was also elevated. We thus further looked at the PERK-ATF4 pathway of the UPR.^26^ Interestingly, while ATF4 typically induces CHOP expression under stress conditions, CHOP did not increase as expected in BA-derived EHCOs, potentially suggestive of abnormal handling with ER stress. A similar phenomenon has been reported in other contexts: Estébanez et al. demonstrated that exercise increases ATF4 without a corresponding CHOP upregulation in elderly individuals.^42^ Konsavage et al. showed that hyperoxia increases both ATF4 and CHOP in adult rats but only ATF4 in neonates.^43^ While PERK-ATF4 pathway was upregulated, AFT6 and IRE1α were not. AFT6 induces antioxidant genes and lack of ATF6 activation leaves cells more vulnerable to oxidative damage.^31^ The downregulation of IRE1α can attenuate its pro-apoptotic functions.^44,45^ Both the lack of ATF6 increase combined with IRE1α decrease may result in cell survival rather than apoptosis of cell with increased ER stress. Downstream genes: *ERO1A, SOD3*, and *WFS1*, regulators of ER stress and antioxidant defense, were significantly downregulated in BA-derived EHCOs. ERO1A knockout has been shown to impair glutathione redox potential in the ER.^46^ Consistent with this, BA EHCOs exhibited decreased GSH levels compared to controls. We previously showed that decreased GSH leads to upregulation of RHOU/Wrch1,andlater downregulation of the transcription factor SOX17^18^ (**Figure 7**).The relationship between ER stress and epithelial polarity has been previously established. ER stress promotes epithelial-mesenchymal transition (EMT) by driving morphological changes, enhancing cell migration, and altering adhesion molecule expression—specifically, downregulating E-cadherin while upregulating N-cadherin and vimentin.^47^ Furthermore, SOD3 has been shown to preserve adherens junction proteins, including E-cadherin^48^ (**Figure 7**). Altogether, our findings suggest that ER stress is not effectively managed in BA cholangiocytes, and this may result in abnormal cell polarity.

An interesting findings of the RNAseq analysis comparing BA patients and controls was differences in drug metabolism and xenobiotics pathways. This is of potential interest as BA is thought to result from an environmental factor be it a toxin, drug or virus during the fetal period in predisposed cholangiocytes.^2,49^ We observed that BA EHCOs demonstrated a more pronounced response to the toxin biliatresone, both by the percentage of morphologically damaged organoids and by the increase in ER stress makers which was higher than that observed in control EHCOs. The ER hosts numerous enzymes, including cytochrome P450 isoforms, that metabolize drugs into more water-soluble forms for excretion. However, in the process of converting xenobiotics, the ER can generate reactive intermediates, free radicals, or other toxic byproducts. These metabolic byproducts can disrupt protein folding, calcium homeostasis, and redox balance within the ER, leading to the accumulation of misfolded or damaged proteins.^50,51^ Our study shows that inhibition of CYP4A activity attenuated ER stress markers, suggesting a crosstalk between xenobiotic metabolism and ER homeostasis (**Figure 7**).

A potential bias of using organoids as human disease model is the potential misrepresentation of the original tissue phenotype under cell culture conditions. This concern is particularly relevant in the study of BA, which is characterized by patchy obstructions along the common bile duct. Our study demonstrated that while most BA-derived EHCOs appear to have an intact structure on light microscopy similar to control-derived EHCOs, gene expression analyses confirmed that BA-derived EHCOs retain distinctive characteristics long after removal from their original environment. Additionally, staining of liver hilum samples obtained from BA patients during Kasai portoenterostomy and non-BA controls obtained from liver explants highlights the involvement of ER stress in BA. Specifically, ER-related proteins were upregulated in the ductal plates of BA patients compared to non-BA controls. These results reinforce the physiological relevance of EHCOs as a model for studying BA pathogenesis. Prior studies have also demonstrated that organoids faithfully maintain the cellular and molecular features of their tissue of origin, both in cholangiocytes^7,52^ and across various other tissues.^53,54^

Taken together, our findings underscore the value of extrahepatic cholangiocyte organoids as a powerful model for studying the pathophysiology of BA. By integrating transcriptomic profiling with functional assays, we reveal key disease-associated alterations, including disruptions in cell polarity, heightened susceptibility to xenobiotic metabolism, and persistent ER stress which is not handled appropriately in BA-derived cholangiocytes. These insights provide a foundation for future research aimed at modulating ER stress responses as a potential strategy to mitigate cholangiocyte injury and improve EHCO function. Importantly, the ability of organoids to retain key pathogenic features of the original tissue highlights their relevance not only for investigating human biliary diseases but also for advancing the development of novel therapeutic interventions.

## Abbreviations

BA: Biliary atresia
BiP: Binding immunoglobulin protein
DE: Differential Expression
ECOs: Extrahepatic cholangiocyte organoids
ER: Endoplasmic reticulum
KPE: Kasai portoenterostomy
RNA-seq: RNA sequencing
TEM: Transmission electron microscopy
UPR: Unfolded protein response

## Acknowledgements

We thank all patients in the cohort for their participation. IGV is a Faculty Fellow of the Edmond J Safra Center for Bioinformatics at Tel Aviv University. KD was supported by the Edmond J Safra Center for Bioinformatics at Tel Aviv University. IGV was supported by European Union Horizon 2020 under grant agreement No. 847422 (ImmunoSep).

We thank the faculty of Medical and Health Sciences at Tel-Aviv University core facility for helping with microscopy imaging.

OWZ was supported by a grant from the Israeli Science Foundation number 1879/21 and by the Fred and Suzanne Biesecker Pediatric Liver Center at the Children’s Hospital of Philadelphia. YH was supported by the Faculty of Medical and Health Science, Tel Aviv University graduate students research grant as part of her PhD studies.

## Supplementary Tables legends

**Table S1**: **Patients’ characteristics**

**Table S2**: **RNA-sequencing differential expression**. For each gene (column 1), its Log2FC, p-value (log10 scaled and signed by direction) and whether it is included in the BA-upregulation or BA-downregulation gene sets are indicated in columns 2,3 and 4, respectively.

**Table S3**: **Differentially expressed pathways between BA and controls**. For each pathway (column 1), reported is its enrichment q-value (calculated with the Wilcoxon test, column 2). Included are only pathways that are enriched with q-value < 0.01. Upregulated and downregulated pathways, reported in column 3, are those that are biased toward positive and negative log2FC scores, respectively. The analysis and FDR correction were applied on all gene sets in the KEGG and GO-Process collections.

**Table S4**: **The effect of perturbations on the BA-upregulated and BA-downregulated genes**. Column 1: A gene, termed also as ‘factor’. Column 2: p-value for the effect of a perturbation in a factor specified in column 1 on the BA-upregulated gene set. Column 3: indication whether the perturbation downregulates or upregulates the BA-upregulated gene set. Column 4: p values for the effect of a perturbation in a factor specified in column 1 on the BA-downregulated gene set. Column 5: indication whether the perturbation downregulates or upregulates the BA-downregulated gene set.

**Table S5**: **Predicted dysfunction of pathways in BA**. Reported are differentially activated pathways (column 1). For each pathway, indicated are the hyper-geometric q-value of this enrichment (column 2) and the category of the factors (column 3). Included are all pathways with q-value < 0.05. The analysis and FDR correction were applied on all gene sets in the REACTOME collection.

**Table S6**: **List of qPCR primers**

**Table S7**: **List of antibodies**

